# Human Beta Oscillations Reflect Magnitude and Fidelity of Priority Shifts in Working Memory

**DOI:** 10.1101/2025.08.07.669112

**Authors:** Nicholas E. Myers, Mark G. Stokes, Paul S. Muhle-Karbe

## Abstract

Flexible prioritisation in working memory (WM) is supported by neural oscillations in frontal and sensory brain areas, but the roles of different oscillations remain poorly understood. Recordings in humans suggest an interplay between prefrontal slow frequency (2-8Hz) and posterior alpha-band (10Hz) oscillations regulating top-down control and retrieval of WM representations, respectively. Complementary work, primarily in non-human primates, suggests an additional role for beta (15-30Hz) oscillations in clearing or inhibiting stimuli from entering WM. Here we investigated the role of neural oscillations in prioritising WM content using electroencephalography (EEG) as participants (humans of any sex) performed a task requiring frequent priority switches between two memorized oriented bars. Behavioural performance revealed switch costs, which scaled with the angular distance between the two items, suggesting that priority shifts are modulated by shift magnitude. Time-frequency analyses revealed increased frontal theta (4–8Hz) and decreased central-parietal beta (15–25Hz) power during switches. Crucially, only beta power scaled with the magnitude of the priority shift and predicted the fidelity of neural decoding of the newly prioritized item during subsequent recall. Theta power, by contrast, was elevated on switch trials but did not vary with update magnitude or decoding strength, suggesting a more general role in signaling control demands. Our findings highlight a particular and previously overlooked role for beta-band oscillations in the flexible prioritisation of WM content.

**Significance Statement:** Working memory permits flexible switching between mental representations, so we can focus on what is most relevant at the moment. Different brain rhythms in frontal control and sensory memory storage areas orchestrate switches but their respective roles remain unclear. Here, using EEG, we show that power reductions of ∼20Hz oscillations over central-parietal regions, usually associated with the motor system, closely track the magnitude of the required switch and the fidelity of the prioritized memory. In contrast, slower 4-8Hz (theta-band) activity over frontal regions increases during priority switches but tracks neither magnitude nor fidelity. Our findings suggest a unique function for central-parietal beta oscillations in the flexible control of working memory.

## Introduction

Working memory (WM) supports the maintenance of task-relevant information, but its capacity is severely limited. Focusing attention on individual items in WM permits effective use of this limited capacity, allowing us to remember more quickly and accurately (Griffin & Nobre, 2003; Souza & Oberauer, 2016; Van Ede & Nobre, 2023). Multiple-state models (Cowan, 2000; Oberauer, 2002) propose that high-priority items are stored in a privileged state that guides perception and action, while other items are concurrently held in a de-prioritised accessory state without driving behaviour (Olivers et al., 2011).

Neurally, prioritised and de-prioritised WM items may be represented in different brain areas (Christophel et al., 2018) and representational formats (El-Gaby et al., 2024; Iamshchinina et al., 2021; Paluch et al., 2025; Panichello & Buschman, 2021; van Loon et al., 2018; Yu et al., 2020). Remarkably, items can be flexibly moved in and out of the focus of attention (Muhle-Karbe et al., 2021; Rerko et al., 2014; Rerko & Oberauer, 2013). Yet the neural mechanisms that underpin priority switches remain to be delineated. This is the focus of the current study.

Prefrontal neural synchronisation for controlled access to items stored in sensory brain areas (Wallis et al., 2015) may represent such a mechanism. More specifically, low-frequency oscillations in the delta and theta frequency ranges (2-8Hz) originating in frontal cortex may play a role in orchestrating top-down control over WM representations stored in sensory or medial temporal areas (Berger et al., 2019; Johnson et al., 2018; Liebe et al., 2012; Roux & Uhlhaas, 2014). Retrieval of these sensory WM representations may occur via spatiotopic desynchronisation of alpha-band oscillations (de Vries et al., 2017, 2018; Myers, Walther, et al., 2015; Poch et al., 2014; Riddle et al., 2020, 2024; van Ede et al., 2016).

Parallel work has emphasised the role of beta oscillations (15-30Hz) in WM (Liljefors et al., 2024; Lundqvist et al., 2016; Miller et al., 2018; Spitzer & Haegens, 2017) but their role in prioritisation is less clear, as priority switches likely require multiple sub-processes that could be linked to beta: a shift in beta synchrony could signal a change in task context (Engel & Fries, 2010) or facilitate removal of the previously prioritised memory (Lundqvist et al., 2018). Given the role of beta in coordinating cortical-basal ganglia loops (Brittain & Brown, 2014; Jenkinson & Brown, 2011), which have been implicated in gating information in and out of WM (Chatham et al., 2014; Chatham & Badre, 2015; D’Ardenne et al., 2012), they may alternatively play a role in selecting a new memory into the prioritised state.

Prioritisation in WM is typically investigated with the retro-cue paradigm, where participants initially encode multiple memory items until a subsequent cue signals which item will be task-relevant for a forthcoming decision (Griffin & Nobre, 2003). This procedure typically incurs only a single priority shift and therefore confounds changes in priority states with changes in memory load, as the initially prioritised item can be dropped after a switch. We recently developed a paradigm that avoids this issue by requiring frequent priority switches between the same two items, so that neither can be dropped from memory. Using this paradigm, we have previously shown that both prioritised and de-prioritised memory items can be decoded from electroencephalography (EEG) recordings and that they track different aspects of memory-guided behaviour (Muhle-Karbe et al., 2021).

Here, we sought to leverage this dataset to characterise the role of theta and beta oscillations during priority switches. We related both oscillations to the magnitude of priority switches and their fidelity by harnessing single-trial decoding of prioritised and de-prioritised memory items. Priority switches increased frontal theta and decreased central-parietal alpha and beta power. We found that beta (but not theta or alpha) oscillations correlated with the magnitude of priority switches and the decoding strength of newly prioritised memory items, but not the previously prioritised item, suggesting that beta oscillations are primarily involved in prioritising new items.

## Materials and Methods

### Participants

We reanalysed data from a previously published study (Muhle-Karbe et al., 2021). 43 adults participated in total. We combined data from two closely related variants of the same visual WM task (experiments 1 and 2). Our previous study showed identical results across both variants. We therefore collapsed all analyses across both experiments. Twenty adults participated in experiment 1 (mean age = 28.1, age range = 18-37, 10 female, 10 male, 1 left-handed, 19 right-handed) and 30 participated in experiment 2 (mean age = 26.8, age range = 19-41, 14 female, 16 male, 2 left-handed, 28 right-handed). Seven participants took part in both experiments, yielding a final sample of 43 participants. For participants who completed both experiments, we analysed data separately within each experiment before averaging the results across the two experiments. All participants reported normal or corrected-to-normal vision and were compensated for their participation. The study was approved by the Central University Research Ethics Committee of the University of Oxford.

### Apparatus

Stimuli were presented using the Psychophysics Toolbox (Brainard, 1997) on a 22” monitor with a refresh rate of 100Hz, viewed at a distance of approximately 60cm in an electrically shielded, sound-attenuated booth with dim background lighting. Stimuli appeared on a grey background ([127 127 127]). Participants responded with left and right index fingers on the “B” and “Y” buttons of a QWERTY keyboard (placement counterbalanced across participants).

### EEG Acquisition

EEG data were collected with a NeuroScan SynAmps RT Amplifier and Curry 7 software. 61 EEG channels were placed following the extended 10-20 positioning system. Channel impedances were kept below 5 kOhm. In both studies, eye movements were recorded via electrooculography (EOG) using electrodes placed above and below the left eye and to the outside of each eye. We also recorded activity in the first dorsal interosseus muscle of the left and right hand via electromyography (EMG). Additional electrodes were placed on the left and right mastoids, which were averaged and used for offline re-referencing. The ground electrode was placed on the left elbow. Data were recorded at 1000Hz.

### Eye-tracking

In some participants (experiment 2, 30 participants) we additionally recorded eye movements using a remote infrared eye-tracker (SR Research, Eyelink 1000) sampling both eyes at 1000Hz. These data were used to confirm that participants maintained close fixation and that EEG-based decoding of memory contents could not be explained by systematic deviations in fixation location (see Muhle-Karbe et al., 2021, for control analyses). Eye-tracking data were not further analysed here.

### Task Design

The experiment used a 2-item visual working memory task requiring frequent priority switches (Figure 1a). Task blocks consisted of 16 trials. At the beginning of each block, participants encoded in WM two orientated bars (6° visual angle in length and 0.25° in width, presented at the same location as subsequent stimuli) in blue ([25.5 25.5 204]) and yellow ([204 204 25.5]). These bars served as memory items for the remainder of the block. The two orientations were drawn randomly from a set of 16 possible orientations (spaced evenly at 11.25° intervals from 2.8125° to 171.5625°) with the constraint that the two items could not be identical or exactly orthogonal. As a result, the absolute angular distance (‘item distance’) between them ranged from 11.25° to 78.75°. One bar colour was associated with a high-pitch tone and the other with a low-pitch tone (mapping counterbalanced across participants), with the two tones serving as retrieval cues for the respective bars later in the block. Participants had unlimited time to encode the two bars and initiated the block via button press.

**Figure 1.**
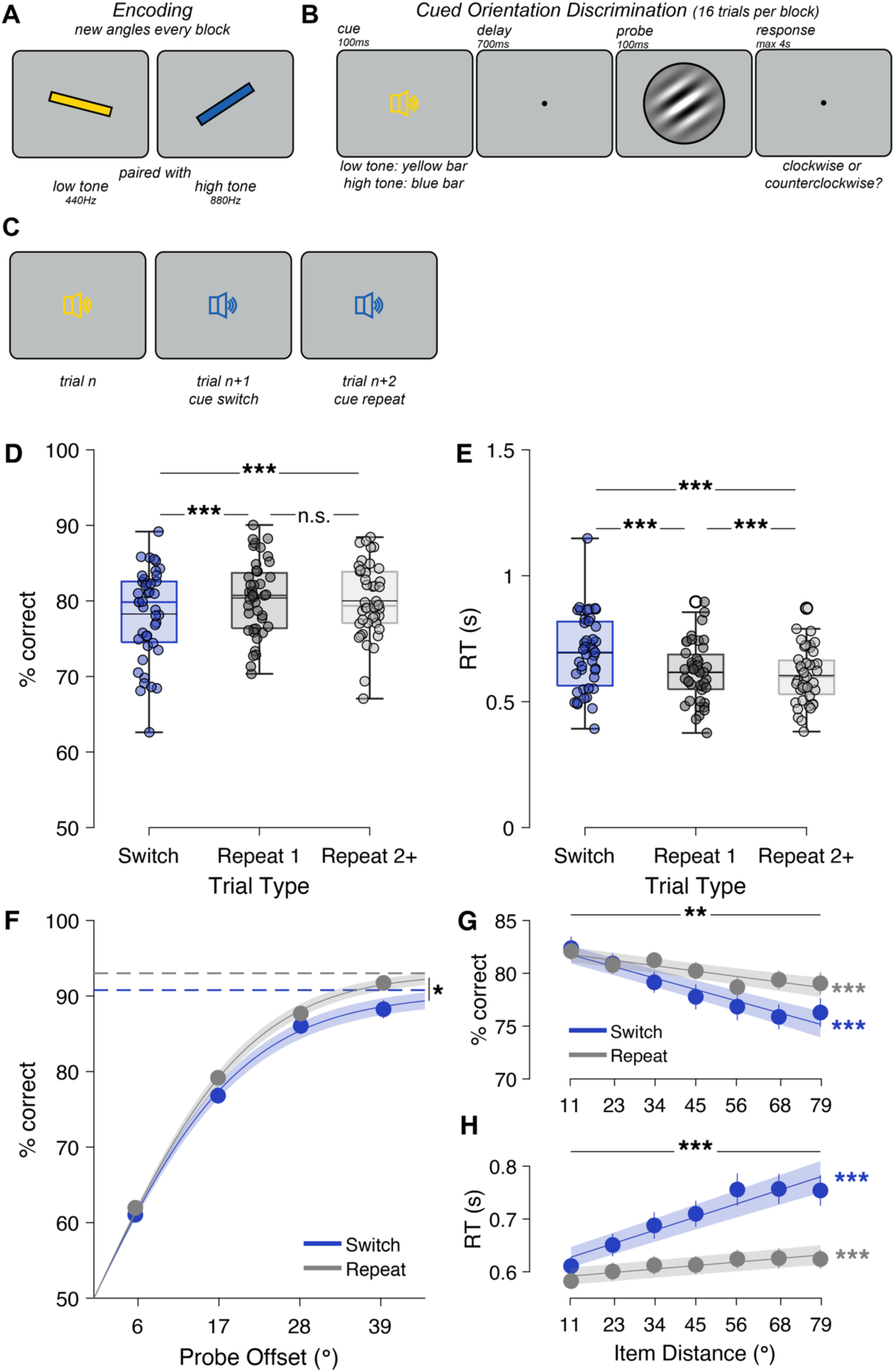
Shifting priority between WM contents incurs a temporary switch cost. **A.** Each block began with encoding of two oriented bars. Each bar was associated with an auditory cue. **B.** On each trial, a cue indicating which WM item was relevant was followed by a probe. Participants judged the probe as clockwise or counterclockwise relative to the cued memory orientation. **C.** Successive trials could cue either the same or different items, requiring frequent priority switches and maintenance of both items throughout the block. **D.** Accuracy was lower on Switch trials than the first Repeat trial after a switch or subsequent Repeat trials (max. 7 repetitions). Coloured horizontal lines show median, black horizontal lines show mean. Dots are individual observers. **E.** Reaction time was slowest on Switch trials and slightly slower on the first Repeat trial. Empty black circles (both Repeat conditions) represent one outlier. **F.** Psychometric curves fit to accuracy had comparable slopes but higher guess rates on Switch trials. Dots are data, with error bars indicating standard error of the mean (s.e.m.). Line is the mean psychometric curve fit (shaded area around mean shows s.e.m.). Horizontal dashed lines are the mean asymptotic performance (‘guess rate’) for each condition (blue: Switch, grey: Repeat). **G.** The similarity between WM orientations (‘Item Distance’) affected accuracy, especially on Switch trials, and accounted for the switch cost. Lines are mean of linear fits to each observer and condition (shaded area shows s.e.m.). **H.** Item distance increased reaction time more on Switch than Repeat trials. ***: p<0.001, **: p<0.01, *: p<0.05, n.s.: not significant.

Within the block, each trial started with a presentation of an auditory cue (pure sinusoidal tones, low tone: 440Hz, high tone: 880Hz, duration: 100ms including a 10ms ramp-up and 10ms ramp-down). The tone signalled which memory item should be used for a forthcoming perceptual decision (cued item), while the other item was maintained for later use in the block (uncued item). The cue was followed by a 700ms delay period within which a black fixation dot was presented centrally on the screen (0.15° diameter). A randomly oriented Gabor patch was presented (orientation drawn from 16 possible orientations, spaced evenly at 11.25° intervals from 8.4375° to 177.1875°; patches had 6° diameter, 50% contrast, 1.75 cycles/°, random phase, and a Gaussian envelope with 1.2° SD). Participants were given a maximum of 4,000ms to judge the probe via button press as clockwise or counter-clockwise relative to the cued memory orientation. Probes were presented for 100ms and replaced by a fixation dot for the remainder of the response period. In experiment 1, the probe was presented centrally on the screen on all trials. In experiment 2, the probe was presented laterally at a distance of 6° from the screen centre. The side of probe presentation (left vs. right) alternated predictably across blocks. A noise patch (Gaussian smoothed random white noise using a kernel with 0.13° SD, convolved with a Gaussian envelope with 1.2° SD) that matched the probe stimulus in luminance, size, contrast, and eccentricity, was presented on the side of the screen at which no probe appeared. Clockwise and counter-clockwise decisions were indicated with the right and left index fingers respectively. The response period was followed by a variable inter-trial interval (400-900ms, drawn from a truncated exponential distribution with mean 550ms).

The next trial cued either the other item (Switch trial) or the same item again (Repeat trial). The likelihood of a cue repeat on each trial was drawn from a modified exponential distribution with a minimum of 1 repeat and a maximum of 7 repeats after each Switch trial. This resulted in comparable trial numbers for Switch trials (464±9, mean±s.e.m.), the first Repeat trial after a switch (Repeat 1, 553±9), and subsequent Repeat trials (Repeats 2-7, abbreviated Repeat 2+, 646±10). Participants received feedback only at the end of each block when their mean accuracy and response time were presented. Accordingly, they had to maintain precise representations of both memory items for the whole duration of the block, as they could not rely on trial-wise feedback to infer the orientation of an item if it was forgotten. Overall, participants completed 128 blocks, resulting in a total of 2048 trials and lasting approximately 2h.

### Behavioural Analysis

We were primarily interested in the effect of priority switches, i.e., trials in which the item cued as relevant by the auditory cue was not the same as the item cued on the preceding trial (Switch trials). We compared Switch trials with Repeat trials, i.e., trials in which the same item was cued as on the preceding trial. We excluded the first trial in each block because it was neither a switch nor a repeat trial. We then calculated mean accuracy and reaction time on switch trials, the first repeat trial (Repeat 1), and any subsequent repeat trials (Repeat 2+). Reaction times were calculated as the median of all correct responses to a given condition. Since accuracy on Repeat 1 and Repeat 2+ trials was largely the same (Fig. 1a), for the remainder of the manuscript we focused on comparing Switch with Repeat 1 trials. Including Repeat 2+ trials in any analyses did not materially affect our results.

We next fit a psychometric curve to the behavioural data to separately estimate the memory precision (inverse of the slope, σ) and the lapse rate (λ) using the response function:

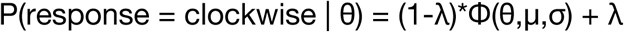

where θ is the angular difference between the probe and cued memory stimulus, λ is the lapse rate, μ is the response bias (fixed to 0), σ is the inverse of the memory precision, and Φ is the cumulative density function of the normal distribution. Precision (1/σ) and lapse rate (λ) were fit using maximum likelihood estimation separately for Switch and Repeat trials. Estimated parameters were compared between conditions using pairwise t-tests.

We extended the psychometric model to include erroneous comparisons to the uncued item (‘swap errors’):

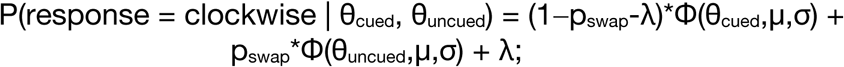

where θ_cued_ is the angular difference of the probe orientation relative to the cued WM item, θ_uncued_ is the angular difference of the probe orientation relative to the uncued WM item, and p_swap_ is the swap rate. To examine the effect of item distance, we fit a linear model of item distance (with values ranging from 11.25°-78.75°) to accuracy and reaction time data on Switch and Repeat trials separately. Regression coefficients were then compared against zero and each other using one-sample or paired-samples t-tests.

### EEG Preprocessing

EEG data were re-referenced to the average of both mastoids. EEG, EOG, and EMG data were down-sampled to 250Hz and bandpass-filtered between 0.1Hz and 45Hz. EEG channels with excessive noise were identified through visual inspection and replaced via interpolation using a weighted average of the surrounding electrodes. The continuous data were epoched from 750ms before to 2500ms after the onset of the cue on each trial. Each trial was inspected visually for blinks, eye-movements, and non-stereotyped artefacts. Trials were rejected if they contained any of those artefacts during the cue-probe delay or probe period. Stereotyped artefacts (from blinks, saccades, or muscle artefacts) outside those periods were subsequently removed via independent component analysis. Unless stated otherwise, the data were baseline-corrected for the decoding analyses using the average signal from the time window of 200 to 50 ms before cue onset.

### EEG Decoding

We conducted multivariate pattern analysis to characterise the neural representations of the cued and uncued WM item. We followed a previously established approach (Muhle-Karbe et al., 2021; Wolff, Jochim, et al., 2020; Wolff, Kandemir, et al., 2020) that used the signal at multiple posterior EEG sensors, pooled across multiple time points, to decode remembered orientations. This approach exploits the dynamic temporal structure of event-related potentials by pooling multivariate information in time, capturing information encoded in spatial activation patterns and in the temporal unfolding of these patterns to achieve greater decoding sensitivity (Wolff, Jochim, et al., 2020).

In keeping with this previous work, decoding analyses were conducted only within posterior EEG sensors (P7, P5, P3, P1, P2, P4, P6, P8, PO7, PO5, PO3, POz, PO4, PO8, O1, Oz, O2). We selected these channels to be consistent with previous studies and because orientation signals are typically expressed most strongly in posterior regions (Cichy et al., 2015; Myers, Rohenkohl, et al., 2015; Wolff, Jochim, et al., 2020). We pooled the signal across a 450ms backward-looking sliding window (i.e., decoding at 800ms after cue onset used data from 350ms-800ms), downsampled to 83.3Hz (i.e., one timepoint every 12ms), yielding 37 timepoints from each of the 17 EEG sensors, or 629 features in total. Data were demeaned across timepoints within each sensor (to reduce any potential carryover of decodable signal from other time windows, although this step seemed to have a minimal effect in our data, see Fig. S5 top vs. bottom row). Our previous work showed that WM decoding peaked around the time of the decision, approximately 200-400ms after probe onset. We therefore focused on the post-probe window for analysis, repeating the decoding analysis in a sliding window from 800-1,400ms relative to cue onset (or 0-600ms relative to probe onset), in 50ms steps.

We computed Mahalanobis distances between the patterns of sensor activity that were evoked by different stimulus orientations and measured the extent to which these distances reflected the underlying circular orientation space. We used a 13-fold cross-validation procedure. On each fold, data from 10 task blocks served as test data and all the remaining data served as training data. From the training data we calculated the average response to each of the 16 orientations, yielding a N_orientations_-by-N_features_ (16x629) training data matrix M_train_. Mahalanobis distances between the 16 average patterns in the training data and each test trial i (M_i_, 1x629) were then computed, using the noise covariance from the training data. Noise covariance was calculated on the residual data after subtracting orientation-specific mean activity from each trial, using a shrinkage estimator (Ledoit & Wolf, 2004). Smaller values indicated a smaller Mahalanobis distance, i.e., greater similarity between the test trial and a given orientation from the training set. For each test trial, this yielded a vector D_i_ of 16 distances. D_i_ was circularly shifted to center the vector on the test trial orientation θ_i_ and inverted so larger values indicated greater pattern similarity, yielding a ‘tuning curve’ that peaks at the centre of the curve when the orientation is decodable. To calculate a trial-wise index of decoding we computed a summary measure that estimated how closely the tuning curve resembled a cosine peaking at the test trial orientation. This index was calculated as the cosine vector mean of the 16-dimensional tuning curve by multiplying each distance with the cosine of the orientation difference to the test trial (Sprague et al., 2016; Wolff et al., 2017):

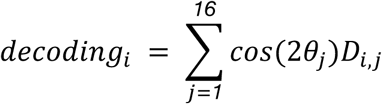

where θ_j_ is the orientation offset of the jth point in the tuning curve (ranging from -78.75° to +90°) and D_i,j_ is the Mahalanobis distance between test trial i and the training set average for the orientation at θ_j_ relative to θ_i_. Angles were doubled in the equation to account for the 180°-symmetry of the orientation stimuli. Because the cosine is zero-centred, if the tuning curve is flat this operation yields a mean decoding score of 0. If decoding peaks at the centre of the curve (at θ_j_=0°), this yields a positive decoding score. An inverted curve (peak near 90°) would yield a negative decoding score. The absolute value of the decoding score is difficult to interpret because it depends on the voltage change over time, the number of sensors, and the number of timepoints. The analysis was run separately for the cued and the uncued orientation. In experiment 2, decoding analyses were conducted separately for blocks with probe presentation on the left and right side, with training and test data matched by side. Training data consisted of all available trials, with decoding accuracy of the test data split afterwards into Switch and Repeat trials.

### EEG Time-Frequency Analysis

Preprocessed EEG data were transformed with a Spatial Laplacian filter to reduce spatial spread of neural sources across the scalp. We used the FieldTrip function *ft_scalpcurrentdensity* (using default conductivity (0.33) and lambda (10^-5^) values and a degree of 10 as in (de Vries et al., 2018)). We then calculated the time-frequency transform in sliding windows from -0.75s before cue onset to 2.5s after cue onset (in 40ms steps). At each timepoint, we calculated power at frequencies between 2 and 40Hz (in 1-Hz steps) using Hanning tapers (using a time window corresponding to 5 cycles of the frequency of interest centered on the current timepoint). We then log-transformed power (10*log10) and baseline-corrected using power from 0.75-0.25s before cue onset.

### Statistical Testing

Unless otherwise noted, we used multiple linear regression to establish relationships between oscillatory power and other task variables (item distance, WM decoding), separately for Switch and Repeat trials. Regressors and dependent variables were z-scored prior to regression. We tested for significance using one-sample t-tests against 0 (at each time point, frequency, or time-frequency combination) and corrected for multiple comparisons via cluster-based permutation testing using 10,000 permutations (Maris & Oostenveld, 2007).

## Results

### WM Priority Switches Incur a Temporary Behavioural Cost

43 Participants performed a visual WM task designed to induce frequent priority switches while we recorded their brain activity through electroencephalographic (EEG) recordings (see Methods for details). The task consisted of 128 mini-blocks each of which contained 16 trials. Participants first encoded two randomly oriented bars into WM and maintained both for the duration of the mini-block. Each bar was associated with a different auditory cue (see Fig. 1a). Every trial began with an auditory cue indicating which of the two items was relevant (Fig. 1b). Participants therefore prioritised the cued stimulus and deprioritised the other (uncued) stimulus. After a 700ms cue-probe interval a randomly oriented probe stimulus appeared and was judged as clockwise or counterclockwise relative to the cued WM orientation. The following trial could then either repeat the same cue (Fig. 1c, Repeat trials, max. 7 repeats in a row) or switch to the other auditory cue (Switch trials), requiring prioritisation of the previously uncued item. Multiple switches occurred in each block, so participants had to keep both stimuli in WM.

Accuracy on Switch trials (78.3%±0.9% s.e.m., Fig. 1d) was modestly but significantly lower than on the first Repeat trial (80.4±0.7%, t_42_=-4.41, p=7.01*10^-5^, Cohen’s d=-0.67) and lower than on subsequent repeat trials (80.0±0.7%, t_42_=-4.06, p=2.08*10^-4^, d=-0.62). The first Repeat trial did not differ from subsequent repeats (t_42_=1.11, p=0.275, d=0.17). Switch costs were driven by an increase in guess rate on Switch trials (9.2±1.1%, Fig. 1f) compared to all Repeat trials (7.0±0.7%, t_42_=2.41, p=0.021, d=0.37). By contrast, precision remained unchanged (Switch: 18.2±1.2°, Repeat: 17.6±0.7°, t_42_=0.43, p=0.67, d=0.07).

Similarly, median reaction time on correct trials was significantly slower on Switch trials (693±23ms, Fig. 1e) compared to either the first Repeat trial (616±18ms, t_42_=9.19, p=1.30*10^-11^, d=1.40) or subsequent Repeat trials (604±16ms, t_42_=9.91, p=1.47*10^-12^, d=1.51). Reaction time on the first Repeat trial was also significantly slower than on subsequent Repeats (t_42_=3.71, p=6.08*10^-4^, d=0.57).

### Priority Switch Costs Scale with Update Magnitude

We next explored whether the similarity between the two items held in WM affected behaviour (Fig. 1g-h). As the angular distance between the cued and uncued item (‘item distance’, ranging from 11-79° in 11.25° steps) increased, accuracy decreased on both Switch trials (decrease in accuracy of 0.97±0.17% per 10 degrees, t-test on slope: t_42_=-5.71, p=1.04*10^-6^, d=-0.87) and Repeat trials (0.46±0.11% per 10 degrees, t_42_=-4.16, p=1.54*10^-4^, d=-0.63). However, the effect of item distance was stronger on Switch trials (t_42_=-3.00, p=0.0045, d=-0.46). After regressing out item distance, there was no longer a significant switch cost (t_42_=0.73, p=0.47, d=0.11), indicating that switch costs were driven primarily by trials requiring a large update from the previous to the currently cued item. Consistent with this, blocks with similar WM item orientations (item distance <25°) showed no switch cost on accuracy (t_42_≤0.29, p≥0.77, d≤0.04).

We found complementary effects on reaction time (Fig. 1e), which increased significantly with item distance on Switch trials (22.5±2.6ms per 10 degrees, t_42_=8.64, p=7.24*10^-11^, d=1.32) and, to a lesser degree, on Repeat trials (5.9±0.9ms per 10 degrees, t_42_=6.60, p=5.4*10^-8^, d=1.01). Again, the effect of item distance was significantly larger on Switch trials (t_42_=7.69, p=1.5*10^-9^, d=1.17). However, even after regressing out the effect a significant difference between Switch and Repeat trials remained (t_42_=2.31, p=0.026, d=0.35). Therefore, reaction time switch costs were present even in blocks with very similar WM items (item distance 11°: t_42_=-3.89, p=3.53*10^-4^, d=0.59).

The item-distance effect could have been driven in part by occasional failures to update the cued item on Switch trials, leading to responses relative to the uncued item (sometimes referred to as a swap error, Bays et al., 2009). When we estimated the swap rate separately from the lapse rate we indeed found that both increased on Switch trials (Suppl. Fig. S1d-e). Nevertheless, when we accounted for potential swap error effects, there was still an independent significant effect of item distance on both accuracy and reaction time (Suppl. Fig. S1f-g).

### Priority Switches Lead to Temporary Decreases in Beta-band Power and Sustained Increases in Theta-band Power

We next examined the neural oscillatory correlates of priority switches in WM. We focused on oscillations because previous work has implicated frontal slow oscillations in the delta/theta-band and posterior alpha-band power reductions (de Vries et al., 2018; Poch et al., 2014; Riddle et al., 2020; Wallis et al., 2015) in switching attention between items held in WM. Here, we observed increased frontal theta power and decreased posterior alpha power, but additionally found a pronounced reduction of power in the beta-band (15-25Hz).

Beta-band power was significantly lower on Switch trials compared to Repeat trials (Fig. 2a). The effect was strongest in a frequency-limited band (approximately 15-25Hz) and peaked in a brief period shortly after the onset of the cue (roughly 400-800ms after cue onset) but before the onset of the memory probe (at 800ms after cue onset). The effect (i.e., decreased power) was most pronounced at central-parietal sites (Fig. 2b) and all subsequent analyses focused on these channels (‘Central Channels’: C1,Cz,C2,CP1,CPz,CP2,P1,Pz,P2). The power decrease on Switch trials at central channels extended to the Alpha band (8-14Hz) and to higher frequencies (25-40Hz, sig. cluster 7-40Hz, p<10^-4^ after cluster-based permutation test) but showed a pronounced peak in the beta-band (Fig. 2c) indicating that beta oscillations are a distinct phenomenon. The effect was significant throughout the trial (from 320ms after cue onset) but strongest in the latter half of the cue period (Fig. 2d).

**Figure 2.**
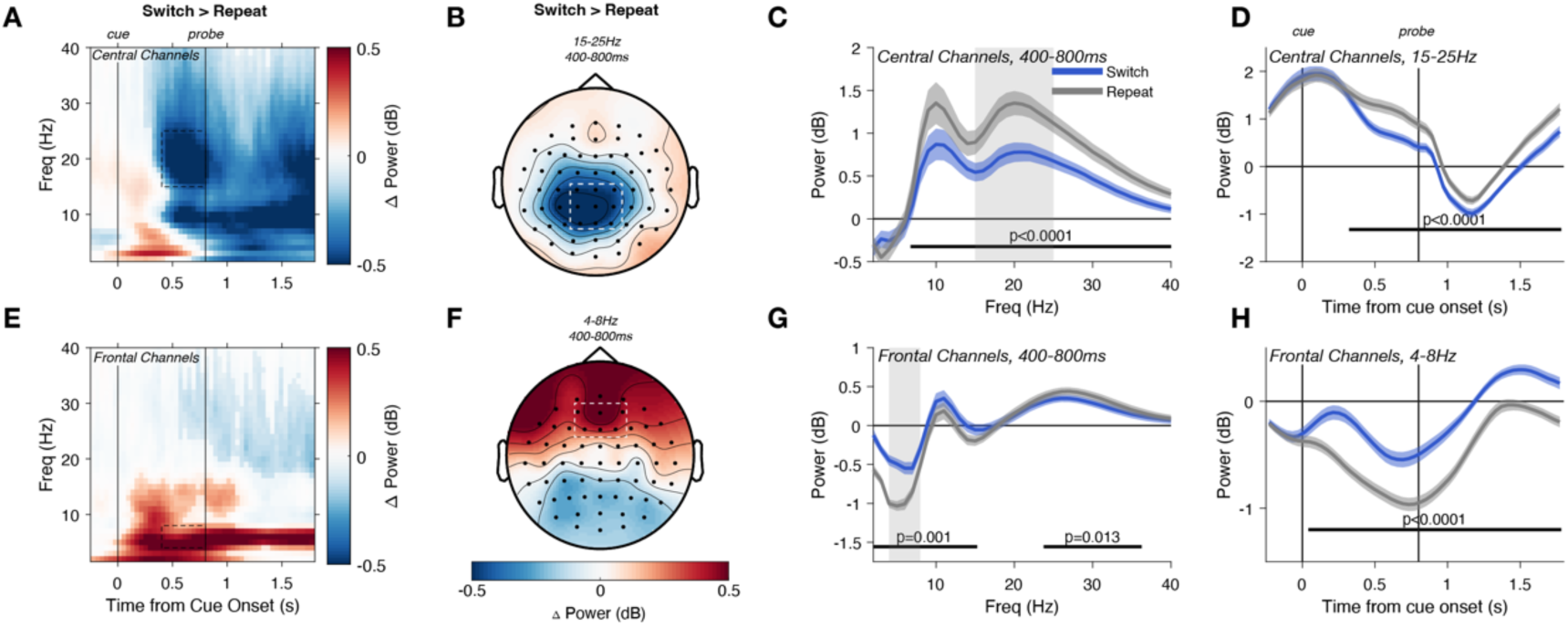
Priority switches led to a temporary decrease in beta-band (15-25Hz) and an increase in theta-band (4-8Hz) power. **A.** Average difference in power between Switch and Repeat trials at central channels. Between the cue and probe onset, power reduction on Switch trials was strongest in the 15-25Hz range but extended from ∼7 to 40Hz (shading indicates significant cluster from permutation test). Time-frequency points are plotted with saturated colours if they belong to a significant cluster (permutation test, p<0.05). **B.** Beta power reduction on Switch trials was strongest at central-parietal channels. **C.** In the late cue period (400-800ms after cue onset), power at central channels peaked in the alpha and beta bands and was lower on Switch trials from 7-40Hz. Shaded area around mean indicates s.e.m., black bar denotes significant difference between Switch and Repeat trials after cluster-based permutation testing. **D.** Central 15-25Hz power was lower throughout Switch trials but strongest in the late cue period. **E-H.** 4-8Hz power at frontal channels was stronger on Switch trials.

In parallel, we observed a significant increase in theta (4-8Hz) power on Switch trials (Fig. 2e) that peaked at frontal midline channels (Fig. 2f, ‘Frontal Channels’: AF3,AFz,AF4,F1,Fz,F2). The effect was limited to the theta and alpha bands (2-15Hz, corrected p=0.001, Fig. 2g) but accompanied by a reduction in higher-frequency power (24-36Hz, p=0.013, Fig. 2g). The increase in theta-band power persisted throughout the Switch trial (Fig. 2h). Finally, power in the alpha-band (8-14Hz) decreased at posterior channels (O1,Oz,O2,PO7,PO3,POz,PO4,PO8, see Supplementary Fig. S2a). We observed the same effects when comparing Switch trials to later Repeat trials (Repeat2+, Fig. S3).

### Beta (but not Theta) Power Scales With Magnitude of Priority Switches

Given the pronounced effect of item distance on behaviour, particularly on Switch trials, we tested if beta- and theta-band effects were also sensitive to item distance. Indeed, beta-band power scaled with item distance (Fig. 3a, F_6,252_=2.303, p=0.035, partial η^2^=0.009), with opposing effects on Switch and Repeat trials (interaction Switch effect with Item Distance effect F_6,252_=8.712, p=1.25*10^-8^, partial η^2^=0.033). On Switch trials, larger priority switches (i.e., larger item distances) led to lower beta-band power (t-test on linear fit, t_42_=-5.74, p=9.54*10^-7^, d=-0.88). On Repeat trials, the effect was reversed, with a (smaller) increase in beta power for larger item distances (t_42_=3.12, p=0.0033, d=0.48), leading to a significant difference in slope between Switch and Repeat trials (t_42_=-5.86, p=6.56*10^-7^, d=-0.89). Even after regressing out the effect of item distance, beta power was still lower on Switch trials (t_42_=-3.42, p=0.0014, d=-0.52). The effect of item distance was confined to the beta band (Fig. 3b, with band-limited sig clusters on Switch trials 15-33Hz, p<10^-4^, on Repeat trials 17-27Hz, p=0.0062, Switch vs. Repeat 15-33Hz, p<10^-4^, Fig. 3e) and to central-parietal channels (Fig. 3c), emerging approximately 200ms after cue onset (Fig. 3d, sig. effect on Switch trials 162-782ms, corrected p<10^-4^, effect on Repeat trials 482-782ms, p=0.0048, difference Switch vs. Repeat 242-782ms, p<10^-4^). Therefore, the beta-band power decrease observed on Switch trials scales with the magnitude of the update between the previous and currently cued item.

**Figure 3.**
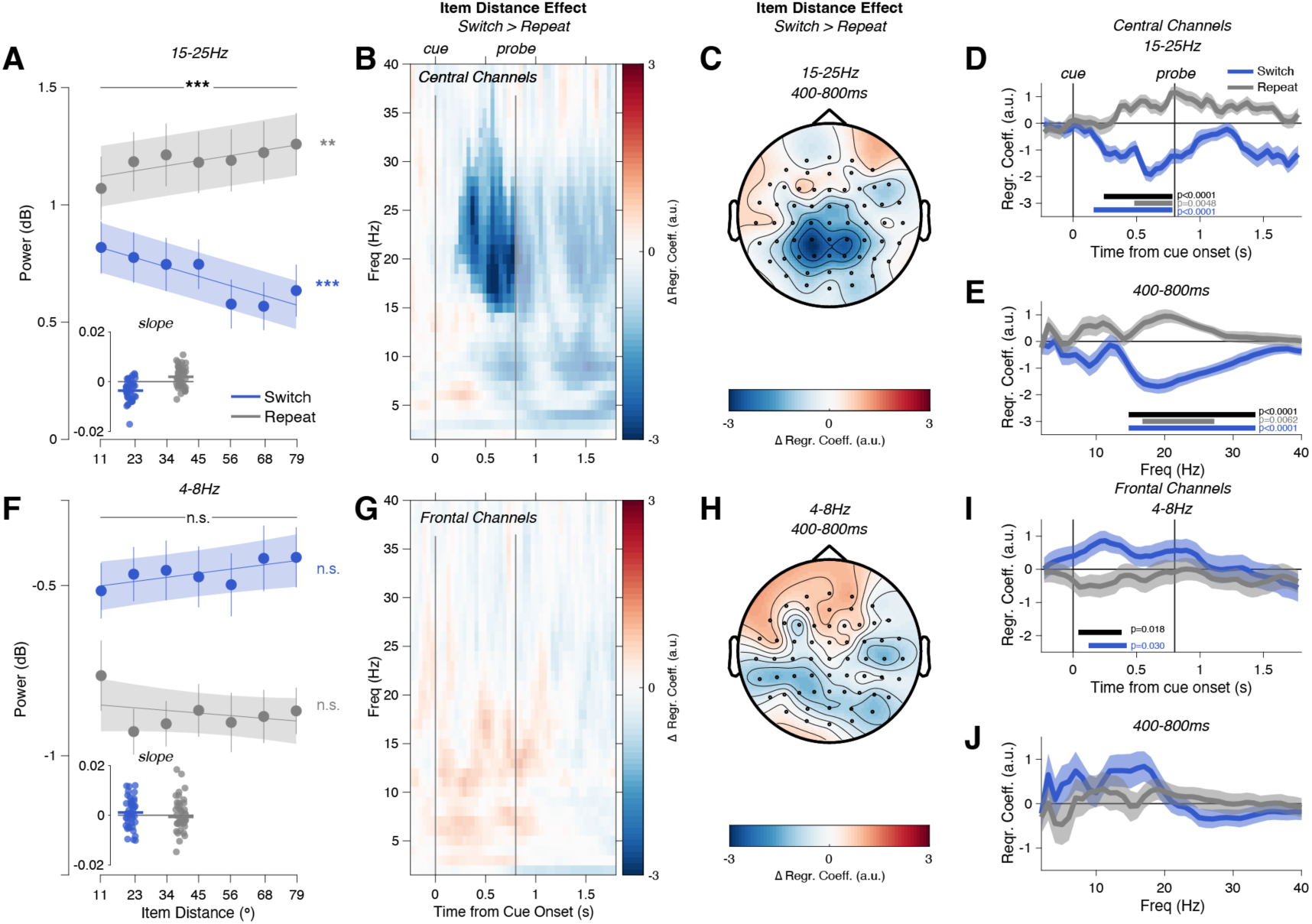
Central Beta Power Scales with Item Distance. **A.** Beta (15-25Hz) power at central channels (400-800ms post-cue), separated by Switch (blue) vs. Repeat (grey) trials and by angular distance between WM items. A large item distance required a larger-magnitude update of the prioritised WM representation on Switch trials. Dots show average power (error bars show s.e.m.), line shows linear fit (shaded area is s.e.m.). Inset shows regression coefficients for each participant. **B.** The effect of item distance on power (regression coefficient on Switches vs. coefficient on Repeats) was resolved over time and frequency and was significant primarily between 15 and 30Hz (area with saturated colour indicates significant cluster from permutation test, p<0.05 corrected). **C.** The effect peaked at central-parietal channels (compare Fig. 2b). **D.** Regression coefficient resolved by time (average of 15-25Hz power at central channels), separately for Switch and Repeat trials. Bars indicate significant effects after correction (blue: Switch vs. 0, grey: Repeat vs. 0, black; Switch vs. Repeat). **E.** Effect resolved by frequency (Central channels, 400-800ms after cue onset). **F-J.** Frontal 4-8Hz power showed no comparable effects, except for a brief correlation on Switch trials immediately after cue onset (panel I).

By contrast, frontal theta power was largely insensitive to item distance (Fig. 3f, main effect of item distance F_6,252_=0.343, p=0.913, partial η^2^=0.001, interaction Switch by item distance F_6,252_=0.867, p=0.519, partial η^2^=0.0034). Linear fits showed no effect of item distance in either Switch (t_42_=1.21, p=0.233, d=0.18) or Repeat trials (t_42_=-0.57, p=0.574, d=-0.09). Theta power remained higher on Switch trials after regressing out item distance (t_42_=3.24, p=0.0024, d=0.49). We found no significant effect at frontal channels when analysed separately by timepoint and frequency (Fig. 3g) or averaged over the late cue period (Fig. 3j). We also saw no clear theta effects at other channels (Fig. 3h). When we examined the effect separately for each time point in the trial, we did find a small effect of item distance on Switch trials immediately following the cue onset (Fig. 3i, effect on Switch trials from 122-422ms, p=0.03, difference between Switch vs. Repeat effects 42-382ms, p=0.018). The early and brief effect suggests that this may reflect a phasic event-related response to the cue rather than the sustained theta oscillation increase observed on Switch trials more generally (Fig. 2e). We found no effect of posterior alpha power on item distance on Switch trials, with only a small effect on Repeat trials (Fig. S2b) that could have been driven by the spatially and spectrally adjacent beta-band effect. In sum, unlike the graded beta-band effect increasing with item distance, oscillatory theta power increases on Switch trials appear to be largely an all-or-none phenomenon.

### Beta Power After Priority Switches Predicts Decoding of Cued WM Item

Since the magnitude of beta power reduction after priority switches scaled with the magnitude of the update, we reasoned that it might be involved in retrieving or shifting attention to the newly cued WM item. We therefore expected that beta power should predict the strength of WM decoding. To this end, we calculated a trial-by-trial orientation decoding score around the time of decision-making after the onset of the probe (200-400ms following probe onset, i.e., 1,000-1,200ms after cue onset). Decoding in this window was maximal (Fig. 4a inset, see also Muhle-Karbe et al., 2021) and comparable on Switch and Repeat trials (Switch trials, significant decoding 800-1,400ms after cue onset, e.g., 0-600ms after probe onset, corrected p<10^-4^, Repeat trials: significant decoding 900-1,400ms, p<10^-4^, no sig. difference between Switch and Repeat, smallest corrected p=0.088).

**Figure 4.**
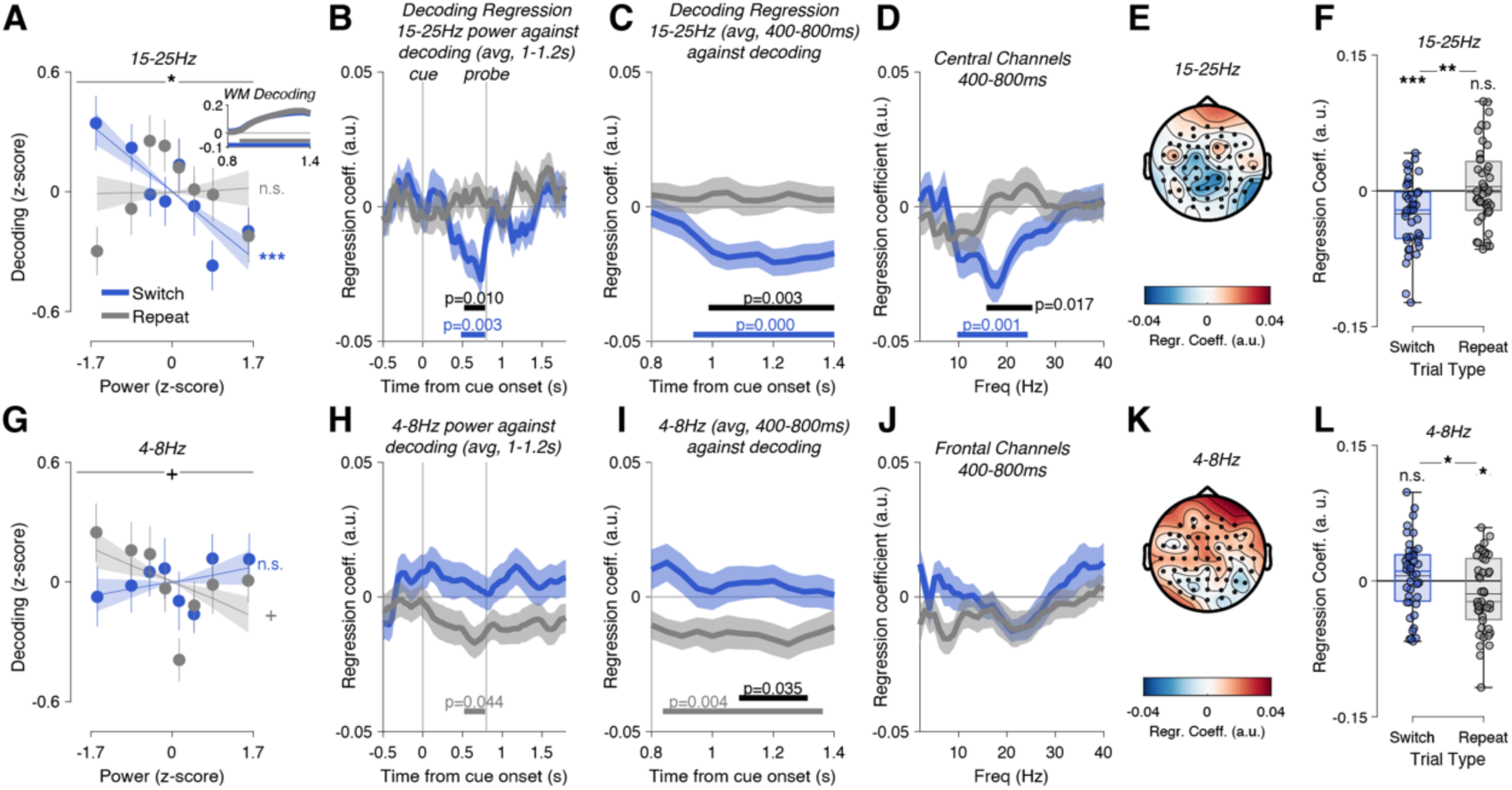
15-25Hz power on priority switch trials predicts decoding of the newly prioritised item. **A.** Switch trials binned by 15-25Hz power (400-800ms, central channels) show a linear relationship with cued orientation decoding. This is not the case on repeat trials. Dots and error bars show mean±s.e.m. of the data. Line and shaded area shows mean±s.e.m. of linear fit. Inset: Timecourse of cued orientation decoding strength (a.u.) from probe onset (0.8s) to 600ms after cue onset (1.4), separately for Switch and Repeat trials. **B.** We regressed beta power across the timecourse of the trial against post-probe decoding (1-1.2s). Only beta power in the late cue interval predicted later decoding (Switch trials only), with no effect on Repeat trials. **C.** Switch trial beta power in the late cue period (400–800ms) predicted decoding throughout the post-probe period, from 150ms after probe onset (0.95s) to the end of the probe epoch (1.4s, 600ms after probe onset). **D.** Cue interval power predicted decoding in a frequency range from 10-24Hz. **E.** Topography of channel-wise power predicting decoding, peaking at central-parietal channels. **F.** Regression coefficient of beta power on decoding from a multiple regression model including item distance and power*item distance interaction. Coloured horizontal lines show median, black horizontal lines show mean. Dots are individual observers. **G-L.** Frontal theta power does not show this effect, but higher theta power predicts lower decoding on repeat trials (but see analysis without baseline correction, Fig. S4).

We then related this to trial-by-trial beta-band power. When we binned all Switch trials by beta power into 8 non-overlapping bins, we saw a clear relationship to decoding: the lower the beta-band power, the higher the decoding (Fig. 4a, linear regression across 8 bins, t_42_=-3.93, p=0.0003, d=-0.60). This was not the case on Repeat trials (t_42_=0.14, p=0.89, d=0.02), leading to a significant difference between slopes (t_42_=-2.68, p=0.011, d=-0.41). Only beta power after the switch cue and shortly before probe onset predicted decoding (Fig. 4b, sig. cluster 492-772ms, corrected p=0.0029, difference to Repeat trials 532-772ms, p=0.0096). Beta power in the pre-probe interval (400-800ms) predicted decoding throughout the post-probe interval (Fig. 4c, 950-1,400ms after cue onset, p=0.0001, difference to repeat trials 1,000-1,400ms, p=0.0033). Decoding before probe onset was much weaker and not correlated with beta power (Fig. S5). At central channels, only alpha- and beta-band power predicted decoding (Fig. 4d,e, 10-24Hz, p=0.001, sig. difference to repeat trials 16-25Hz, p=0.0172).

Because of the relationship between item distance and both beta power and accuracy, these effects could have been driven by differences in item distance. To account for this possibility, we conducted a multiple regression analysis with (z-scored) beta power, z-scored item distance and their interaction as predictors and WM decoding as the dependent variable. The effect of beta power on decoding remained significant for Switch trials (Fig. 4f, t_42_=-4.420, p=6.82*10^-5^, d=-0.67) and non-significant for Repeat trials (t_42_=0.672, p=0.506, Switch vs. Repeat: t_42_=3.428, p=0.0014, d=-0.52) and we saw no difference in the effect of item distance on decoding between Switch and Repeat trials (t_42_=1.03, p=0.31 for model including 15-25Hz power, t_42_=0.79, p=0.43 for model including 4-8Hz power), and there was no significant interaction between beta power and item distance on decoding (p>0.22).

We saw no comparable effect of theta power on Switch trials. In fact, there appeared to be only a modest effect of theta power on decoding on Repeat trials. This effect trended into an unexpected direction, with higher theta power predicting lower decoding (Fig. 4g,h, sig. cluster 532-772ms, p=0.044, also after correcting for item distance, Fig. 4l). This effect occurred in a similar time range (Fig. 4i) as the beta power effect but appeared to be statistically weaker and showed no clear peak in the frequency spectrum (Fig. 4j) or strong effects at non-frontal channels (Fig. 4k). More importantly, it appeared to depend on the choice of baseline and may have been driven by fluctuations in pre-trial theta power. When we conducted the analysis without baseline-correction (Fig. S4b) or after regressing baseline power out of the signal (Fig. S4d), there was no longer a significant relationship between theta power and decoding. Instead, increased theta power on switch trials predicted increased decoding of the cued item late in the trial, mirroring the beta power effect (Fig. S4b,d, third panel). Similarly, alpha power had no effect on decoding of the cued item with or without baseline correction (Fig. S2c,f). The beta power effect (Fig. S4a,c), on the other hand, remained unchanged and significant.

In sum, beta power had a significant effect on decoding only on switch trials, suggesting a role in prioritising the newly relevant WM item. Conversely, an *insufficient* reduction in beta power might reflect insufficient or ineffective deprioritisation of the uncued item and thus predict its decoding remaining high. To test this possibility, we repeated the analysis but with trial-by-trial decoding of the uncued item. Decoding of uncued orientations was highly significant on both Switch (1-1.4s after cue onset, corrected p=0.0001) and Repeat trials (1.05-1.4s, p=0.0028, Switch>Repeat 1.25-1.4s, p=0.0295, Fig. 5a, left panel inset). Nevertheless, beta power had no effect on decoding (main effect: F_6.29,264.14_=0.72, p=0.64, Switch trials: t_42_=0.66, p=0.51, Repeat trials: t_42_=0.78, p=0.44, Fig. 5a). This remained true after regressing out the effect of item distance (all p>0.51, Fig. 5a, right panel). Theta power also did not predict decoding of the uncued item (main effect: F_5.91,248.24_=0.319, p=0.92, Switch trials: t_42_=-1.06, p=0.30, Repeat trials: t_42_=-0.56, p=0.58, Fig. 5b). Regressing out the effect of item distance did not change this result (all p>0.20).

**Figure 5.**
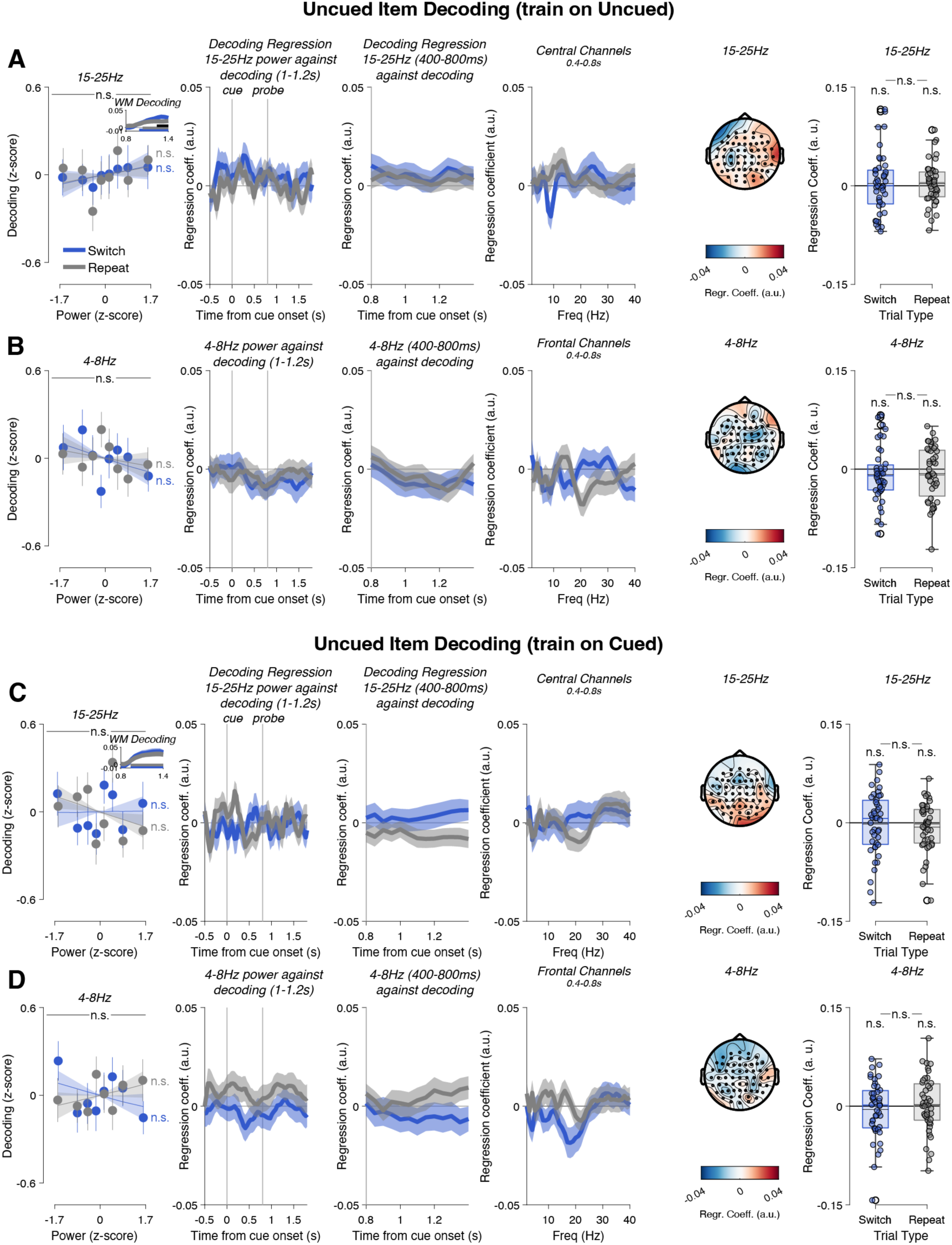
15-25Hz and 4-8Hz power on priority switch trials do not predict decoding of the uncued item. **A.** No relationship between 15-25Hz power and decoding of the uncued item. **B.** Similarly, 4-8Hz power had no effect. **C-D.** Crossdecoding the uncued item (train on cued, test on uncued) also showed no effect. Conventions as in Fig. 4.

A related possibility is that insufficient beta desynchronisation on Switches leads to lingering activation of the uncued item in the representational format of a cued item (i.e., in a prioritised state). To test this, we trained a decoder on cued WM items and tested it on uncued items. We again found no significant effect. Cross-decoding of uncued orientations (training on cued orientations) was significant on both Switch (0.95-1.4s after cue onset, corrected p=0.0001) and Repeat trials (0.95-1.4s, p=0.0002, Switch>Repeat n.s. Fig. 5c, left panel inset). Beta power had no effect on decoding (main effect: F_6.15,258.32_=1.78, p=0.10, Switch trials: t_42_=0.05, p=0.96, Repeat trials: t_42_=-1.26, p=0.21, Fig. 5c). This remained true after regressing out the effect of item distance (all p>0.25, Fig. 5c, right panel). Theta power also did not predict decoding of uncued orientations (main effect: F_6.14,258.03_=0.418, p=0.87, Switch trials: t_42_=-0.95, p=0.35, Repeat trials: t_42_=0.80, p=0.43, Fig. 5d). Regressing out the effect of item distance did not change this result (all p>0.41, Fig. 5d, right panel). Alpha power again had no effect on decoding of the uncued item (Fig. S2d,e).

We conclude that beta power reduction affects only the activation of a newly cued item but not the removal of a previously relevant item from the prioritised representational state.

## Discussion

Our study examined the role of neural oscillations in priority switches in working memory. Previous studies had identified predominantly theta and alpha oscillations, with relatively less emphasis on beta-band oscillations, but their respective contributions have remained unclear. Here we identified a central role for transient reductions in beta-band power in two aspects of switching priority in working memory: the magnitude of the representational shift and the fidelity of the update, as measured by the decoding strength of the newly prioritised item. By contrast, while priority switches also affected theta and alpha power, neither of those oscillations strongly correlated with update magnitude or fidelity.

At the behavioural level, priority switches incurred a temporary cost that manifested in reduced accuracy and increased reaction time, consistent with classic work on task switching (Koch et al., 2018; Monsell, 2003). The reduction in accuracy was linked to an increase in guess rate but no change in precision, suggesting that switch costs may have arisen because of occasional failures to prioritise but not because the representation itself was noisier. In line with this interpretation, switches also increased the rate of swap errors. Interestingly, the cost of switching priorities between memories scaled linearly with the angular distance between the two items. This was a surprisingly strong effect that explained the entirety of the switch cost on accuracy (although swap errors may have contributed to part of this effect). The dependence of switch costs on memory similarity suggests that updating the focus of attention is a graded operation, akin to mental rotation (Shepard & Metzler, 1971). This finding appears to be at odds with models of WM as a storage system for arbitrary content (Bocincova et al., 2022; Bouchacourt & Buschman, 2019; Manohar et al., 2019) since it suggests that control operations in WM reflect the representational space of features in WM. A comparable similarity effect has recently been reported for switch costs between different task rules (Bustos et al., 2024). Cognitive control operations more generally may therefore depend on the representational geometry of task-relevant features, consistent with reports of map-like representations for other cognitive functions (Bellmund et al., 2018).

At the neural level, we observed a robust beta-band (approx. 15-25Hz) power reduction following switch cues, particularly at central-parietal electrodes. Power reduction scaled with item distance, suggesting that beta oscillations reflect the magnitude of the required switch. Notably, this effect was specific to Switch trials and absent (or modestly reversed) on Repeat trials, while behaviour on Repeat trials still showed an effect of item distance. This specificity to switch trials suggests a role for beta in the switching process, rather than coding for item distance per se. It is unclear why switches between more dissimilar items require a greater reduction in beta amplitude. One possibility is that beta tracks the duration of the switching process, and switching to a more dissimilar item takes longer (as with mental rotation). More dissimilar Items could also be encoded in more spatially distant populations, possibly requiring stronger beta reduction (Lundqvist et al., 2023) during a switch. We stress, however, that these are speculative explanations that cannot be resolved with the current data.

Importantly, beta power on priority switches also predicted the fidelity of the updated representation. On Switch trials, lower beta power was associated with stronger decoding of the newly prioritised item later in the decision phase. The correlation was specific to beta frequencies, to the cued (but not the uncued) item, and occurred only on Switch trials. This suggests that beta oscillations are involved in the selection and prioritisation of relevant memory items rather than the removal of de-prioritised items. This conclusion appears to contrast with previous reports of beta oscillations playing a key role in clearing items from WM when they are no longer needed (Lundqvist et al., 2018). In our paradigm, however, items were not fully removed from memory, as uncued items could still be probed later in the block. Beta oscillations may therefore play distinct roles in updating and clearing WM (Lundqvist et al., 2023, 2024). Beta-band desynchronisation has previously been implicated in increasing the coding capacity of neural populations (Hanslmayr et al., 2012) to aid reinstatement of long-term memories (Griffiths et al., 2019), consistent with the proposed role in updating in our task. We also observed a temporal progression of events, with beta desynchronisation occurring before probe onset but affecting decoding after probe onset. While it is possible that this was simply because stronger decoding after the probe made it easier to find a significant correlation, it may hint at a temporal sequence in which a reduction in beta power initiates a preparatory cascade of events from identifying and accessing the cued item to then prioritising it before this is reflected in improved decoding when the cued item is needed for decision-making. We speculate that reductions in beta bursts may not only serve to prepare for upcoming external updates of WM (Liljefors et al., 2024) but also for internal updates within the focus of attention. Consistent with this view, a related line of work has suggested that beta synchronisation helps in the selection and downstream routing of behaviourally relevant neural ensembles (Rassi et al., 2023; Salazar et al., 2012; Spitzer & Haegens, 2017).

While the limited spatial resolution of EEG prevents us from drawing strong conclusions about anatomical sources of the beta-band effect, its well-documented involvement in regulating activity between cortex and the basal ganglia (Brittain & Brown, 2014; Engel & Fries, 2010; Jenkinson & Brown, 2011; Schmidt et al., 2019) suggests beta synchrony in corticostriatal loops could orchestrate WM prioritisation. Along these lines, the basal ganglia have been proposed as a source of selecting from WM for the next action (Chatham et al., 2014; Hazy et al., 2006). Other work has suggested a direct role for beta desynchronisation in the motor system in WM tasks which permit preparation of an action before recall (Boettcher et al., 2021; Schneider et al., 2017). This is unlikely in our case because participants could not anticipate the correct response (probe rotated clockwise/counterclockwise) at the time of the peak beta effect (before probe onset). More broadly, surprise at the less frequent Switch trials (approx. 30% of trials) could have led to inhibition of the motor system (Wessel & Aron, 2013, 2017), though it is unclear why such an effect would scale with update magnitude or predict the strength of decoding. In sum, while several processes likely occur during prioritisation, the specificity of the beta-band effects in our study are most consistent with a role in selecting and prioritising new information.

Theta oscillations, while elevated on Switch trials, did not scale with item distance and showed no consistent relationship with decoding strength. This suggests that theta may reflect a more general, possibly all-or-none, process signaling control demands rather than the implementation of specific updates. This interpretation is consistent with the apparent role of delta/theta in regulating posterior alpha desynchronisation when space-based selection of WM contents is possible (de Vries et al., 2020). It also aligns with a longstanding proposal that midline frontal theta signals cognitive control demands more generally (Cavanagh & Frank, 2014) rather than executing control.

Alpha oscillations, like beta, showed a significant reduction on Switch trials in our study. However, as with theta, we did not observe any correlations with update magnitude or decoding. Spatially lateralised alpha oscillations have previously been linked to successful prioritisation of cued WM contents (Boettcher et al., 2021; Myers, Walther, et al., 2015; Poch et al., 2014; Wallis et al., 2015). This discrepancy may be due to the non-spatial nature of our task, where items were cued via auditory tones rather than by spatial location. Future work could untangle the contributions of different prioritisation mechanisms by comparing spatial with non-spatial selection.

Taken together, our results highlight a previously underappreciated role for beta oscillations in the dynamic reorganisation of WM contents. Beta power reduction appears to facilitate the transition of items into the prioritised state, with the degree of reduction reflecting both the magnitude of the update and the quality of the resulting representation. We saw no such relationship with the uncued item, indicating that beta in our task was not directly correlated with deprioritising previously cued information. However, swap errors were more frequent after priority switches, indicating occasional failures to update the focus of attention, and beta could be relevant to this process or in suppressing the uncued item more generally. While recent work has begun to address the neural mechanisms of swap errors in working memory (Alleman et al., 2024), the role of neural oscillations in this process is unknown and appears to be a fruitful avenue for further research. These findings more broadly highlight the potential relationship between neural oscillations and interference-free maintenance of multiple task representations (Fang et al., 2025). Future work could aim to further dissociate the causal role and specific subprocesses of priority switching supported by beta-band oscillations by examining prioritisation efficacy when beta oscillations are entrained artificially or when they are affected by neurological conditions such as Parkinson’s disease.

## Acknowledgments

This work was supported by funding from the Wellcome Trust (Award 201409/Z/16/Z/ to N.E.M. and Award 210849/Z/18/Z to P.S.M.K), University College, Oxford (N.E.M.), and Linacre College, Oxford (P.S.M.K.), the Biotechnology and Biological Sciences Research Council (Award BB/M010732/1 to M.G.S.), the James S. McDonnell Foundation (Award 220020405 to M.G.S.), and the National Institute for Health Research Oxford Health Biomedical Research Centre. The Wellcome Centre for Integrative Neuroimaging is supported by core funding from the Wellcome Trust (Grant 203139/Z/16/Z). This article is dedicated to Mark G. Stokes, who passed away in 2023.

## Supplementary Results

**Figure S1.**
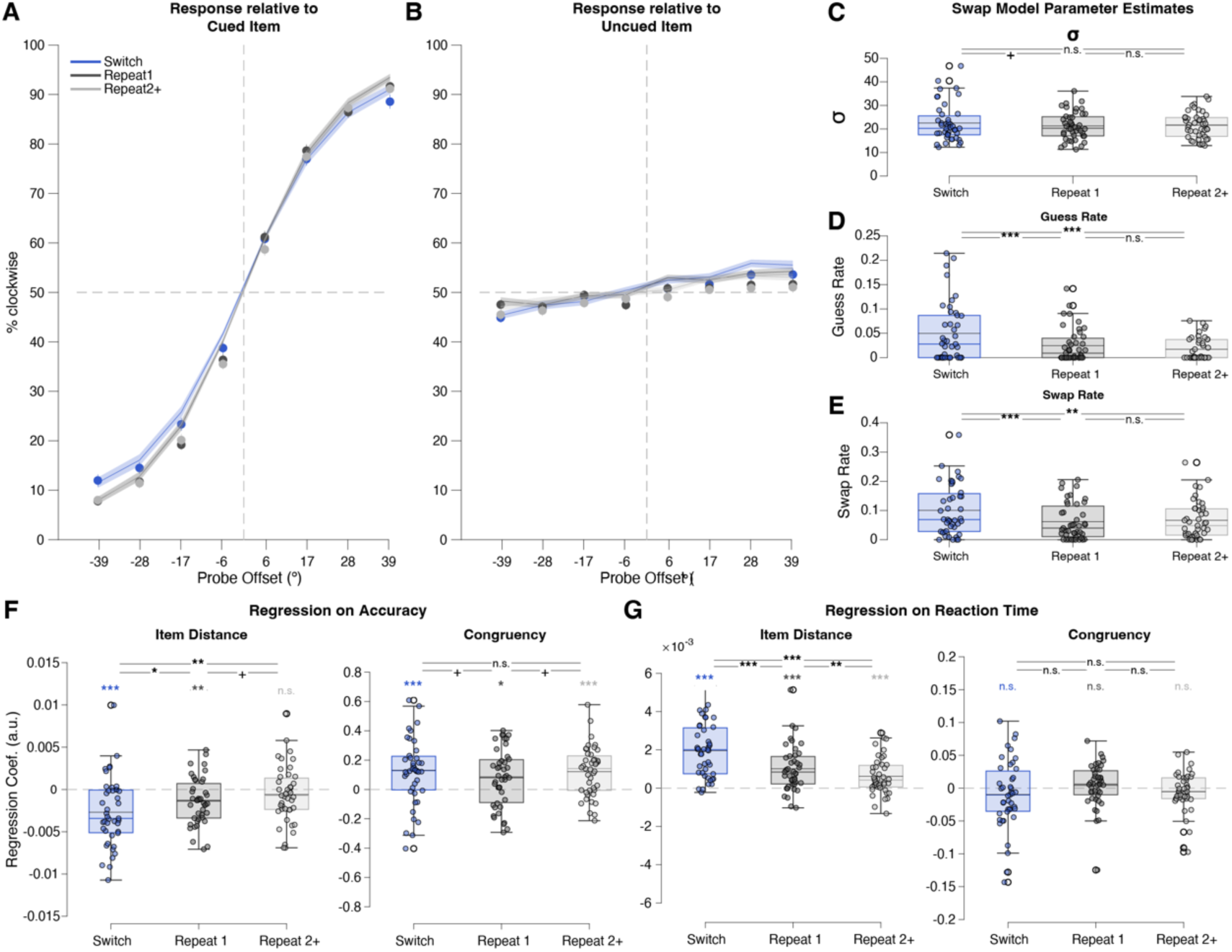
Independent Effects of Swap Errors and Item Distace on Switch trials. **A.** Psychometric curves showing proportion of trials judged clockwise, sorted by probe orientation relative to cued WM item. **B.** Responses sorted relative to the uncued item, indicating swap errors. Dots show data (whiskers are standard error of the mean), lines show predicted response from fit of a mixture model (shaded area is standard error the mean across participants). **C.-E.** Mixture model parameters (inverse precision, guess/lapse rate, and swap rate) are shown in the right panels. **F.** Effect of item distance (left panel) and congruency (right panel) on accuracy. **G.** Effect on reaction time.

Inverse precision (σ), p_guess_, and p_swap_ were fit using maximum likelihood estimation separately for Switch and Repeat trials.

Swap errors were significantly higher on Switch trials (Switch: 10.1±1.3%) than on first Repeat trials (Repeat1: 6.2±0.9%, t_42_=3.99, p=2.566*10^-4^, Cohen’s d=0.61) or subsequent Repeat trials (Repeat2+: 6.6±1.0%, t_42_=3.16, p=2.930*10^-3^, d=0.48). Swap errors did not differ between Repeat1 and Repeat2+ trials (t_42_=-0.73, p=0.469, d=-0.11).

Precision still did not differ significantly between Switch (22.5±1.2°) and Repeat1 (21.3±0.9°) trials (t_42_=1.84, p=0.073, d=0.28), although there was a trend. Precision on Repeat2+ trials (21.6±0.8°) was no different from Switch trials (t_42_=1.38, p=0.174, d=0.21) or Repeat1 trials (t_42_=-0.64, p=0.526, d=-0.10).

Similarly, the guess/lapse rate was still significantly higher on Switch (5.0±0.9%) than on Repeat1 (2.5±0.5%) trials (t_42_=3.66, p=6.987*10^-4^, d=0.56) and Repeat2+ trials (1.7±0.4%, t_42_=3.61, p=8.15*10^-4^, d=0.55). Guess rates did not differ between Repeat1 and Repeat2+ trials (t_42_=1.42, p=0.164, d=0.22).

While the mixture modeling suggests that swap errors contributed to switch costs, it does not rule out that item distance contributed as well. We quantified both effects in tandem using multiple regression. We reasoned that if swap errors contributed to behaviour, they would reduce accuracy when the correct response relative to the uncued WM item was different from that to the cued item (i.e., incongruent trials). For example, if the cued item was 30° and the probe orientation was 45°, the correct response would be ‘counterclockwise’. If the uncued item was 60°, a swap error might produce a ‘clockwise’ judgment, resulting in an incorrect response. By contrast, swap errors should not reduce accuracy on congruent trials. We used trialwise logistic regression to predict accuracy from item distance and congruency.

We also included a number of nuisance regressors (the absolute angle between the probe and the cued item, between the probe and the uncued item, the block number, and the trial number within the block) which we had found to influence behaviour in our previous publication (Muhle-Karbe et al., 2021). We then compared regression coefficients against 0 and across Switch vs. Repeat trials. We repeated the same analysis (using linear regression) on reaction time (on correct trials only).

Even after taking potential swap errors into account (as measured by congruency), item distance had a significant negative effect on accuracy on Switch trials (t_42_=-4.36, p=8.2*10^-5^, Cohen’s d=-0.66) and on the first Repeat trial (t_42_=-3.29, p=2.04*10^-3^, d=-0.50), but not subsequent trials (t_42_=-1.182, p=0.244, d=-0.18). The effect on Switch trials was marginally stronger compared to the first Repeat trial (t_42_=2.03, p=0.0486, d=-0.31) and compared to subsequent Repeat trials (t_42_=3.35, p=0.00171, d=-0.51). The effect was not different on the first vs. subsequent Repeat trials (t_42_=-1.82, p=0.0756, d=-0.28), although there was a trend.

At the same time, congruency also affected accuracy on all three trial types (Switch: t_42_=3.65, p=7.25*10^-4^, d=0.56; Repeat1: t_42_=2.6427, p=0.0115, d=0.40; Repeat2+: t_42_=4.67, p=3.05*10^-5^, d=0.71). The congruency effect was marginally stronger on Switch trials than the first Repeat trial (t_42_=-2.001, p=0.0519, d=0.31), consistent with the psychometric modeling results above, but not on Switch vs. subsequent Repeat trials (t_42_=-0.345, p=0.732, d=0.05). The effect was also marginally but non-significantly smaller on the first Repeat trial compared to subsequent Repeats (t_42_=-1.837, p=0.0733, d=-0.28).

In sum, item distance and congruency (as a measure of swap errors) made independent contributions to performance, and both appeared to be modulated on Switch trials.

For completeness, we also examined the effect of item distance and congruency on reaction time (RT). In brief, item distance had a significant effect on RT that was strongest on Switch trials, while congruency had no effect on RT.

Item Distance had a significant slowing effect on RT on all trial types (Switch: t_42_=9.12, p=1.62*10^-11^, d=1.39; Repeat1: t_42_=5.57, p=1.65*10^-6^, d=0.85; Repeat2+: t_42_=4.27, p=1.11*10^-4^, d=0.65). The effect was stronger on Switch trials than on the first (t_42_=-5.89, p=5.71*10^-7^, d=0.90) or subsequent Repeats (t_42_=-8.35, p=1.83*10^-10^, d=1.27). The effect was also stronger on the first Repeat compared to subsequent Repeat trials (t_42_=2.84, p=7.00*10^-3^, d=0.43).

Congruency had no significant effect on RT on any trial type (Switch: t_42_=-1.21, p=0.232; Repeat1: t_42_=0.85, p=0.399; Repeat2+: t_42_=-1.14, p=0.261). There were no differences between conditions (Switch vs. Repeat1: t_42_=1.59, p=0.118, d=-0.24; Switch vs Repeat2+: t_42_=0.55, p=0.582, d=-0.08; Repeat1 vs Repeat2+: t_42_=1.51, p=0.137, d=0.23).

**Figure S2.**
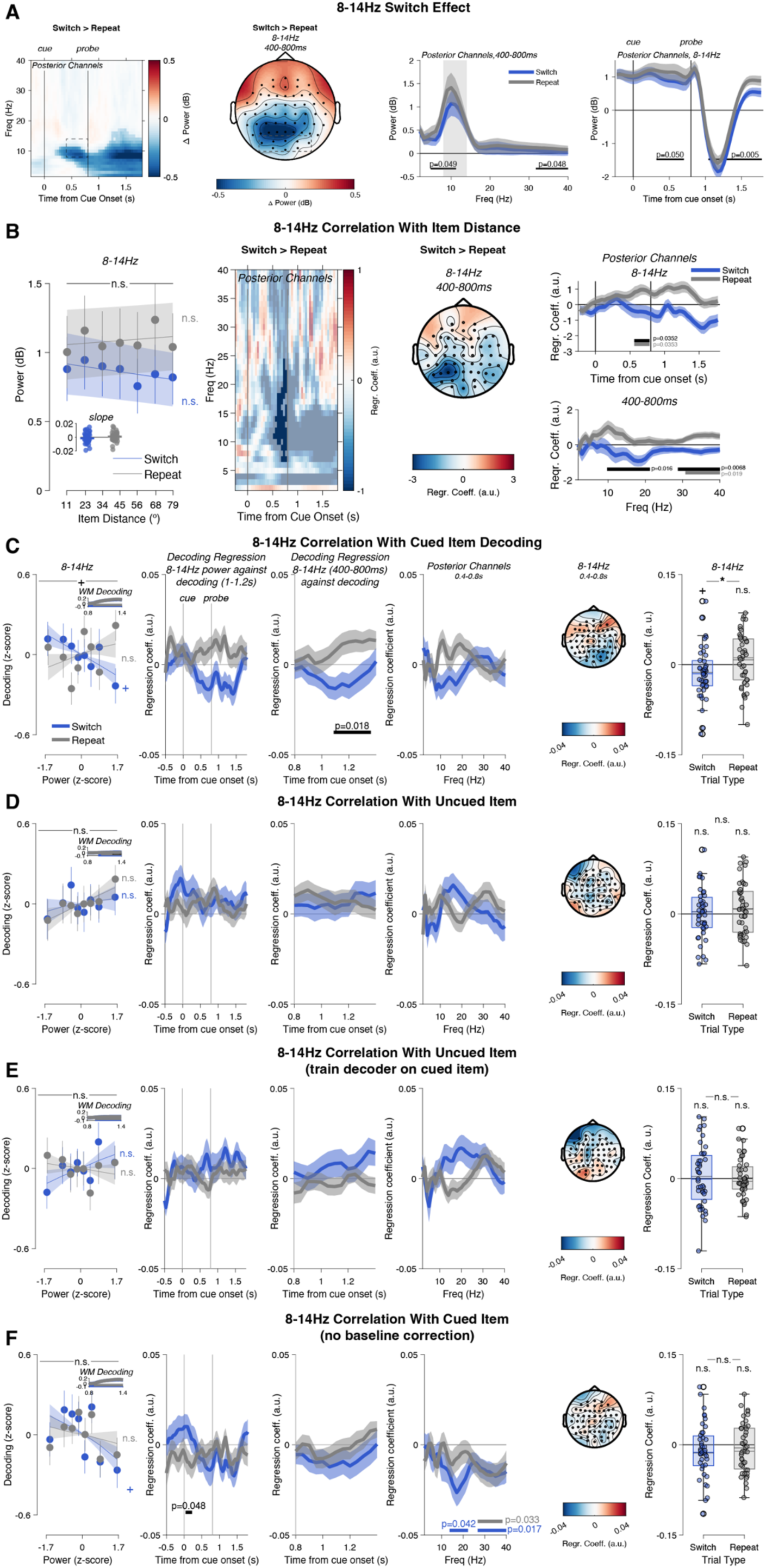
**A.** Posterior Alpha Power desynchronised on Switch trials. Desynchronisation was most pronounced in the low alpha band (<10Hz) on Switch trials (left panel). Desynchronisation was strongest at posterior-parietal sensors (second panel). The desynchronisation appears limited to the theta-low alpha range. We also observed a desynchronisation between 30-40Hz on Switch trials (third panel). The timecourse of desynchronisation is limited to a short period in the second half of the cue-target interval (right panel). Plotting conventions as in Fig. 2. **B.** Posterior Alpha Power does not scale with item distance on Switch trials. There is, however, a trend towards a positive relationship between alpha power and item distance on Repeat trials. Plotting conventions as in Fig. 3. **C.** No relationship between alpha-band desynchronisation and decoding of the cued item, **D.** uncued item, or **E.** cross-decoding of the uncued item (trained on the cued item). **F.** There is also no effect when alpha-band power is used without baseline-correction. Plotting conventions as in Fig. 4.

### Supplementary Results of the Analysis of Item Distance Effects on Alpha Power

Alpha power in the late cue period (400-800ms, 8-14Hz, posterior channels) was lower on Switch trials (*F_1,42_=5.80, p=0.021,* partial η^2^*=0.12*) but did not scale with item distance (*F_6,252_=1.41, p=0.21, partial* η^2^*=0.006,* left panel).

There was also no interaction between Switch and item distance (*F_6,252_=1.162, p=0.33, partial* η^2^*=0.004*). A linear regression of item distance on alpha power did not show any effect on Switch trials (*t_42_=-1.25, p=0.22, d=-0.19*) or Repeat trials (*t_42_=0.88, p=0.38, d=0.13*). Nevertheless, when examining the difference in regression slopes between Switch and Repeat trials separately for each timepoint and frequency, we found a small but significant cluster (p=0.0234) late in the cue-target interval that extended into the beta band (approximately 500-800ms, 7-24Hz, second panel). This effect was restricted to posterior sensors (third panel) and appeared to be driven by a small positive effect on Repeat trials late in the delay (Repeat>0, 562-782ms, p=0.0353, Repeat>Switch, p=0.0352, upper right panel). The effect was not restricted to the alpha band, ranging from 10-21Hz (p=0.016). It thus may have depended to some extent on the beta-band effect (Fig. 3). An additional effect appeared in the low gamma band (31-40Hz, p=0.019).

**Figure S3.**
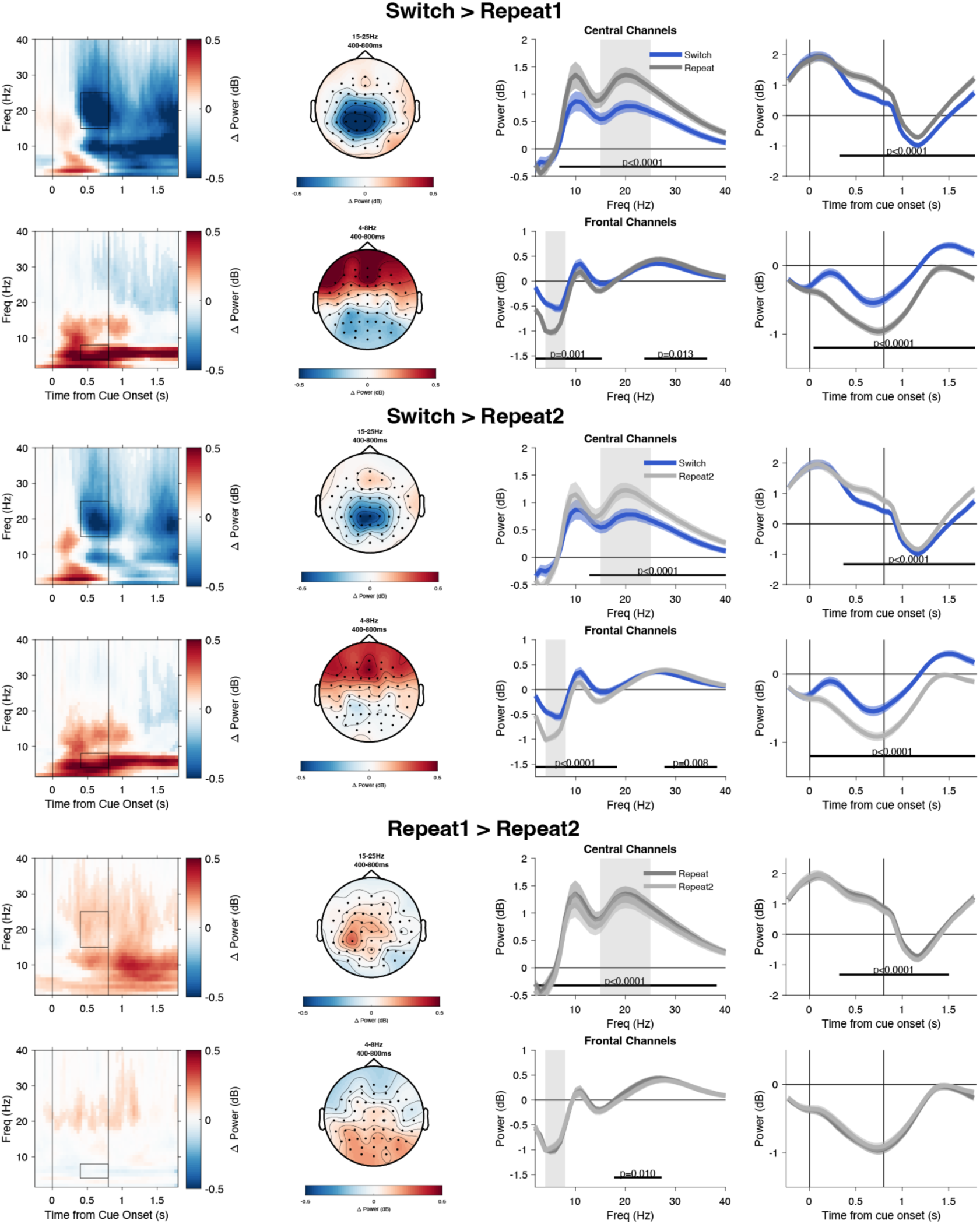
Beta- and Theta-power differences between Switch trials, first Repeats (Repeat1), and subsequent Repeats (Repeat2). Switch trials differed from Repeat1 trials (rows 1-2, from Fig. 2, repeated here for comparison) and from Repeat 2 trials (rows 3-4). While Repeat1 and Repeat2 trials were qualitatively similar, Repeat1 trials showed lower central alpha/beta power. There were no theta-band differences. Plotting conventions as in Figure 2.

**Figure S4.**
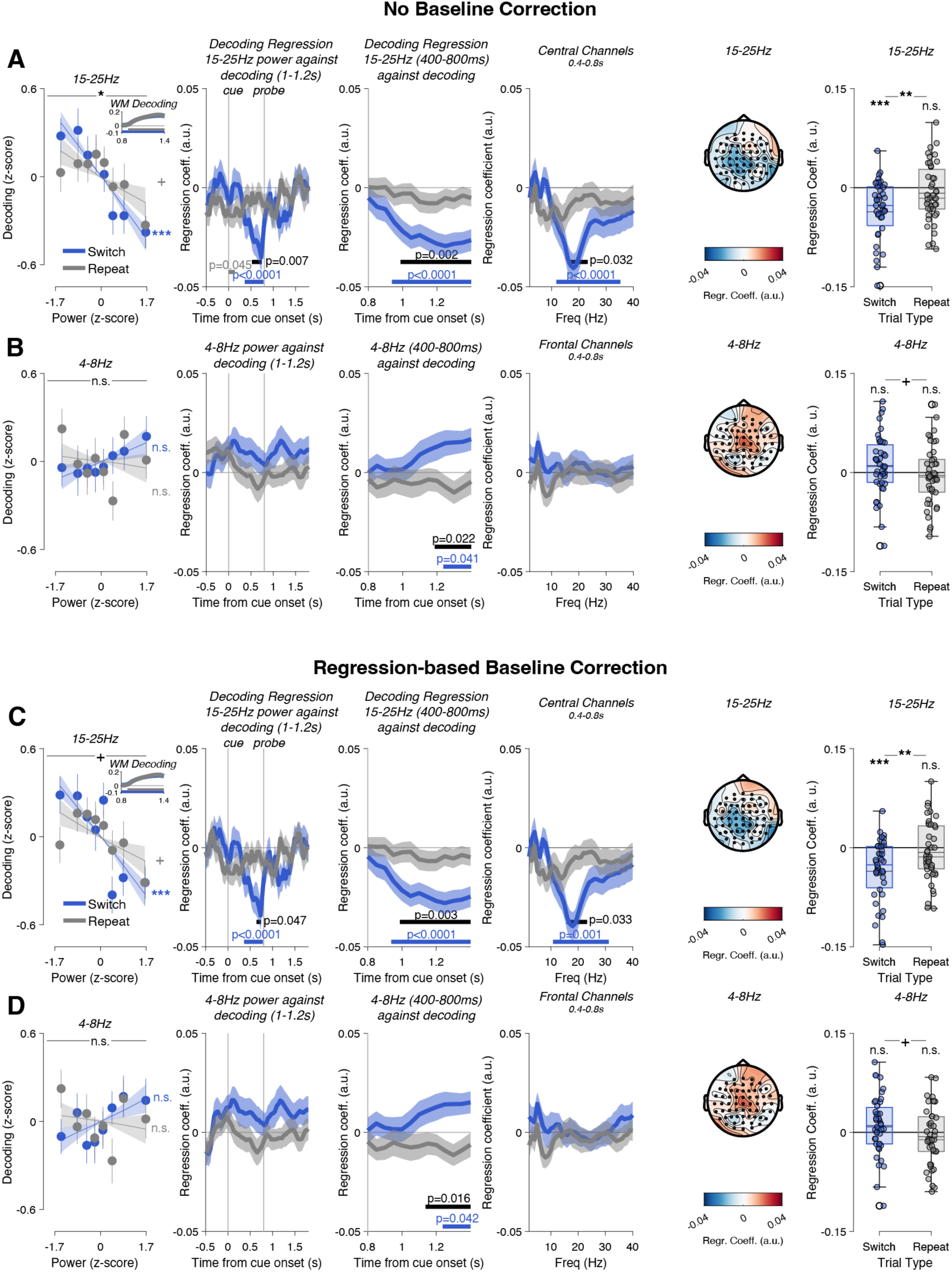
**A.** Omitting baseline correction has no effect on the relationship between beta-power and decoding on Switch trials. **B.** However, it substantially affects the relationship between theta-power and decoding, where now a positive correlation emerges for decoding late in the trial. **C.** Alternatively, regressing baseline power out of the signal also has no effect on the relationship between beta-power and decoding on Switch trials. **D.** Again it affects the relationship between theta-power and decoding, where a positive correlation emerges for decoding late in the trial. +, p<0.10. Other plotting conventions as in Figure 4.

**Figure S5.**
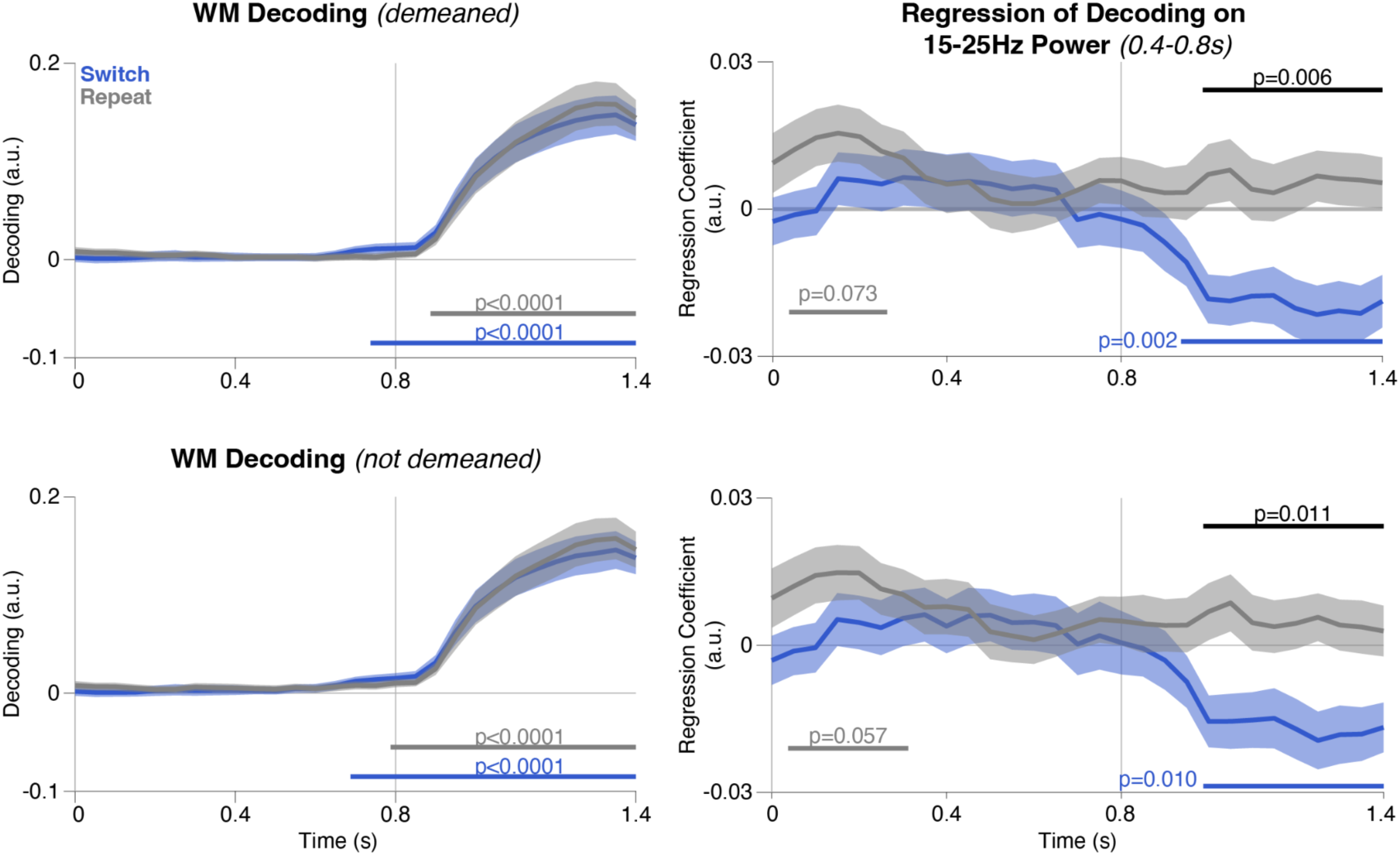
Decoding emerges around the time of probe onset (0.8s), but is much weaker in the cue-probe interval (left panels). Cued item decoding is indistinguishable on Switch (blue) and Repeat (grey) trials. Beta power on Switch trials (0.4-0.8s) only predicted decoding after probe onset (right panels). When EEG features used for decoding were not demeaned over time (bottom row), the results remained virtually unchanged.

